# The neonicotinoid insecticide imidacloprid disrupts bumblebee foraging rhythms and sleep

**DOI:** 10.1101/2020.04.07.030023

**Authors:** Kiah Tasman, Sean A. Rands, James J.L. Hodge

## Abstract

Neonicotinoids have been implicated in the large declines observed in flying insects such as bumblebees, an important group of pollinators[1]. Neonicotinoids are agonists of nicotinic acetylcholine receptors that are found throughout the insect central nervous system, and are the main mediators of synaptic neurotransmission[2]. These receptors are important for the function of the insect central clock and circadian rhythms[3, 4]. The clock allows pollinators to coincide their activity with the availability of floral resources, favourable flight temperatures, as well as impacting learning, navigation and communication[5]. Here we show that exposure to the field relevant concentration of 10 µg/L of imidacloprid can cause a reduction in foraging activity and reduce both locomotor and foraging rhythmicity in *Bombus terrestris*. Foragers showed an increase in daytime sleep and an increase in the proportion of activity occurring at night. This would likely negatively impact foraging and pollination opportunities, reducing the ability of the colony to grow and reproduce, endangering crop yields.

## INTRODUCTION

Bumblebees are a diverse and important group of pollinators, being major pollinators of both crops and flowers. Of the five most important crop pollinators in Europe, three are bumblebees[6]. Many crops are particularly reliant on bumblebee pollination[7, 8], and crop pollination in Europe is worth over €22bn *per annum* and is essential for food security[6]. Unfortunately, despite their ecological and economic value, bumblebees face dramatic population declines, with 46% of species in Europe in decline and 24% threatened with extinction[6]. Due to crop lost to insect pests, demand for insecticides remains high[9, 10]. The most common insecticides worldwide are neonicotinoids, global sales of which are worth US$1 billion/year[2, 10]. Neonicotinoids have the same mechanism of action, being agonists of nicotinic acetylcholine receptors (nAChR), the main neurotransmitter system in the insect nervous system, and share target site cross-resistance[11]. They were branded safe compared to their predecessors because they do not act on mammalian nAChRs[2, 11]. However, before their introduction to market, safety tests were not adequately performed on beneficial insects, for which neonicotinoids have proven potent neurotoxins.

Most research on the effects of neonicotinoids on pollinators has used the honeybee, *Apis mellifera*[12]. However, honeybees and bumblebees show differential responses to neonicotinoids, with bumblebees appearing more susceptible to lethal and sublethal effects[13]. This high-lights the importance of increasing the diversity of pollinators studied to determine the ecological consequences of neonicotinoid use. Concentrations as low as 1 µg/L (or 1 part per billion (ppb)) of imidacloprid, can cause reduced foraging motivation in *Bombus terrestris*[14]. A dose of 6 ppb of imidacloprid can disrupt bumblebee nest behaviour[15, 16] and large field studies have shown decreased bumblebee populations across the EU as a result of neonicotinoid exposure[17]. Neonicotinoids have high solubility and persistence in the environment[18], meaning insects are still at risk of exposure, despite the current EU ban on imidacloprid.

Due to the abundance and importance of nAChRs in the insect central nervous system, the potential sub-lethal effects of neonicotinoids are very broad and their effect on many vital behaviours of pollinators such as circadian rhythms and sleep are still unknown. The circadian clock is integral to pollinator foraging efficiency as flower opening, scent release and nectar production are dependent on time of day[5]. Circadian rhythms also affect other behaviours, such as caring for offspring and learning, with honeybees learning novel, rewarding odours better in the morning[5, 19]. This helps them find new foraging patches, due to most flowers being nectar-rich in the morning[5]. The bee anatomical and molecular clock closely resemble that of *Drosophila*[20], in which circadian entrainment, synchronicity within the clock and communication between the light sensing organs and the central clock are reliant upon nAChR signalling[21-24], as are post-synaptic mushroom body (MB) output neurons which regulate sleep[25, 26]. *Bombus terrestris* also possess similarly grouped and located clock neurons to *Drosophila*, with conserved clock genes such as *period (per)*, neuropeptides such as pigment dispersing factor (PDF) and neurotransmitters such as ACh[5, 35-37].

In the honeybee the clock neurons project to brain regions including visual circuitry, pars intercerebralis and pars lateralis[38], both of which control locomotor activity and sleep in *Drosophila*[38-41]. PDF and nAChR expressing lateral pacemaker clock neurons with extensive branching patterns are also present in other insects, including crickets and cockroaches[3, 20, 27, 28], demonstrating the clock circuitry is conserved across insects. Honeybee clock neurons are important for the sun-compass pathway[29], a vital navigational tool in bees allowing communication of resource location *via* the waggle dance in honeybees[30]. The clock also dictates the timing of sleep, which is required for memory consolidation and synaptic homeostasis[31-36]. Furthermore, sleep timing in bumblebees is important for round the clock care of offspring[37]. Therefore, circadian and sleep disruption is likely to have detrimental effects to the pollination services, behaviour and fitness of beneficial insects. Due to the importance of nAChRs signalling in the insect clock and sleep centres, we hypothesised neonicotinoids would disrupt bee rhythmicity and sleep. We therefore tested the effect of field relevant concentration of imidacloprid on *Bombus terrestris* foragers.

## RESULTS

The activity of isolated *Bombus terrestris* foragers in individual tubes was measured using the locomotor activity monitor to assess behaviour in both 12hrs:12hrs light:dark (LD) conditions and constant darkness (DD). Rhythmicity under LD conditions was studied as this provides a more naturalistic reflection of how day/night behaviour may be affected in the field and also allows sleep to be investigated. The removal of light cues under constant conditions in DD reveals the endogenous circadian rhythm and clock function. Exposure to field relevant concentrations of 1-10 µg/L imidacloprid disrupted the rhythmicity and quantity of locomotor activity of isolated foragers (Figure 1A). The rhythmicity statistic (RS) was calculated as a measure of rhythm strength, with RS>1.5 by convention taken to indicate rhythmic behaviour[38]. In LD conditions, imidacloprid decreased mean rhythmicity (Figure 1C) and both 1 and 10 µg/L increased the proportion of foragers that were arrhythmic (Figure 1B), from 10% in control foragers to 36% and 67% respectively. Imidacloprid also reduced the total activity of foragers, with 1 µg/L reducing activity during both day and night, and 10 µg/L reducing daytime activity (Figure 1D).

**Figure 1:**
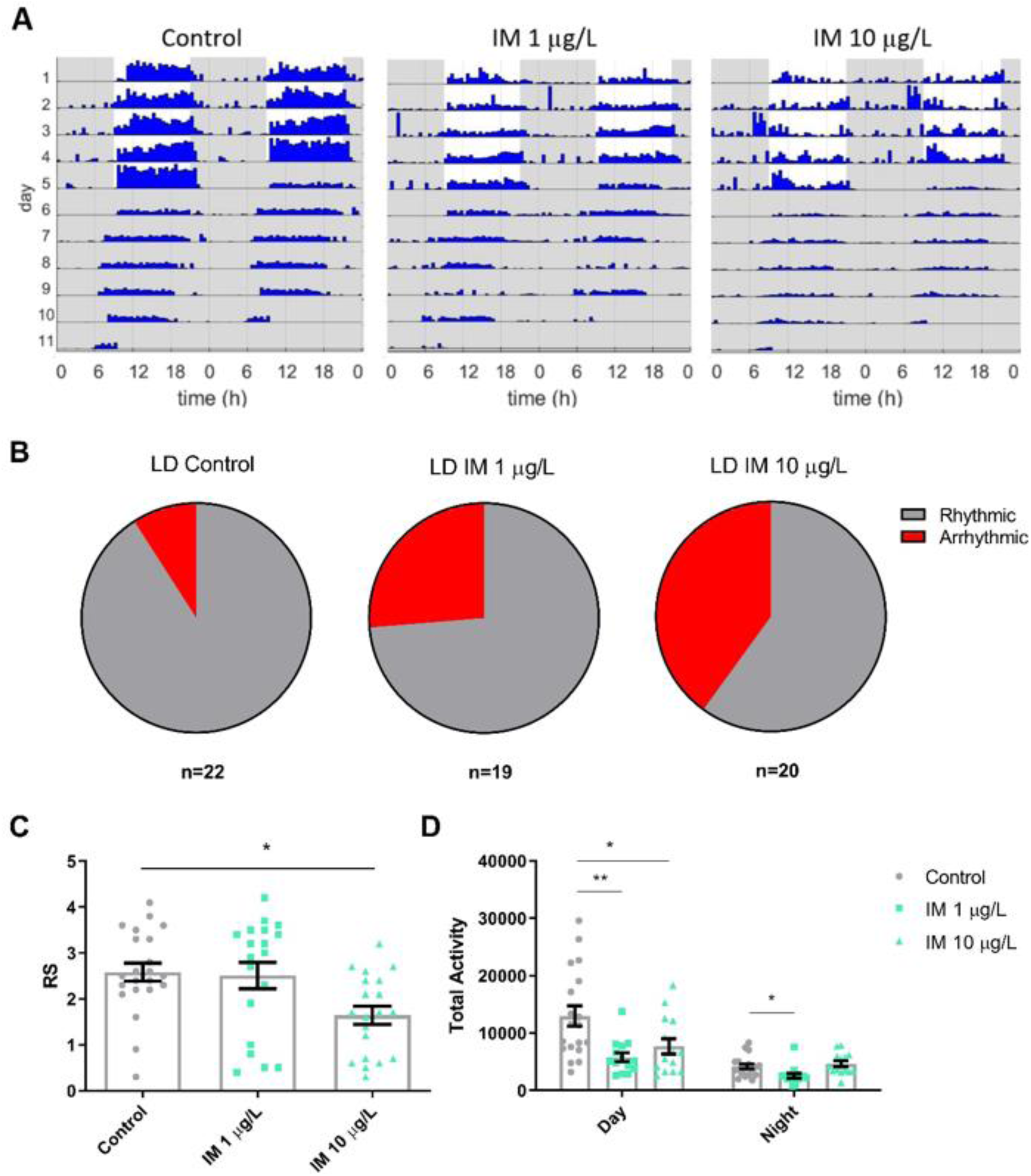
Field relevant concentrations of imidacloprid affects rhythmicity in isolated bumblebee foragers in 12hr:12hr light:dark conditions. (A) Representative actograms for a *Bombus terrestris* forager on control food, or food containing 1 µg/L or 10 µg/L imidacloprid (IM). (B) Proportion of foragers that were arrhythmic (rhythmicity statistic (RS)≤1.5) in LD for each treatment. (C) Mean rhythmicity for either control foragers or those fed 1 or 10 µg/L imidacloprid, in LD conditions (*F*_2,58_=5.3, p=0.008). (D) Mean locomotor activity for foragers in each treatment group in LD conditions, during the day (*F*_2,44_=6.7, p=0.003) and the night (*F*_2,44_=5.4, p=0.008). Each data point represents a single bee, n=19-22 bees for each treatment group.

Conversely, in DD, exposure to either concentration of imidacloprid had little effect on the forager’s activity or rhythmicity (Figure 2). Foragers fed 1 or 10 µg/L imidacloprid had the same mean rhythmicity and levels of activity as control foragers (Figure 2B-C). The proportion of each population that were arrhythmic was also similar, with 40% of control foragers arrhythmic compared to 33% at 1 µg/L imidacloprid and 50% at 10 µg/L (Figure 2A).

**Figure 2:**
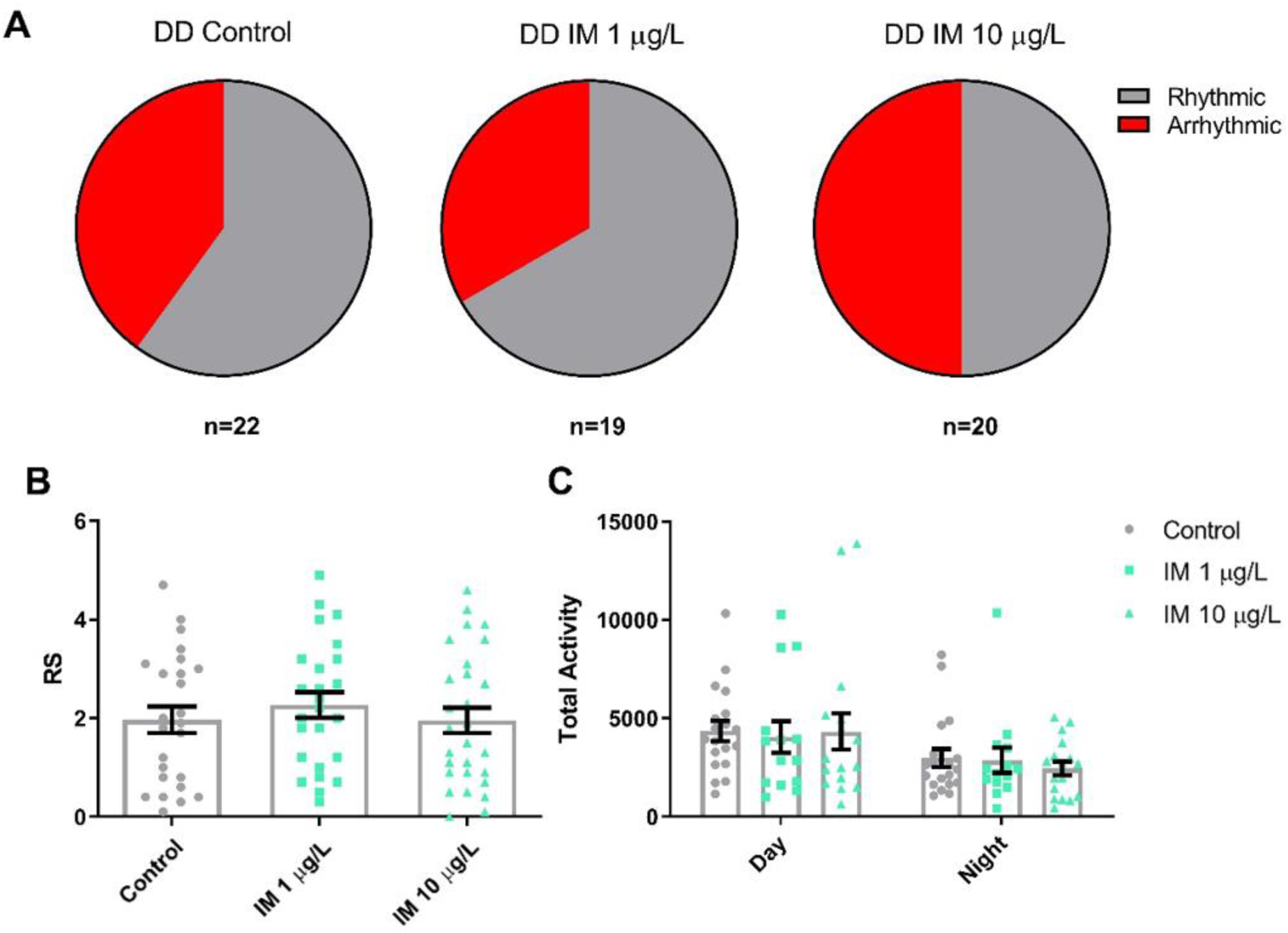
Field relevant concentrations of imidacloprid do not reduce rhythmicity in isolated bumblebee foragers in constant darkness. (A) Proportion of *Bombus terrestris* foragers that were arrhythmic (RS≤1.5) in DD for foragers on control food or food containing 1 or 10 µg/L imidacloprid (IM). (B) Mean rhythmicity (Rhythmicity Statistic (RS)) for foragers in each treatment group in DD conditions (*F*_2,47_=0.5, p=0.637). (C) Mean locomotor activity for foragers in each treatment group in DD conditions, during the subjective day (*F*_2,47_=0.1, p=0.947) and night (*F*_2,47_=0.6, p=0.541). Each data point in the histograms represents a single bee, n=14-22 bees for each treatment.

Foragers who were fed 10 µg/L imidacloprid showed an increase in sleep compared to controls, particularly during the day (Figure 3A-B). This is likely due to the increased number of daytime sleep episodes (Figure 3C) initiated by these foragers. The length of these sleep episodes was the same as in control foragers (Figure 3D).

**Figure 3:**
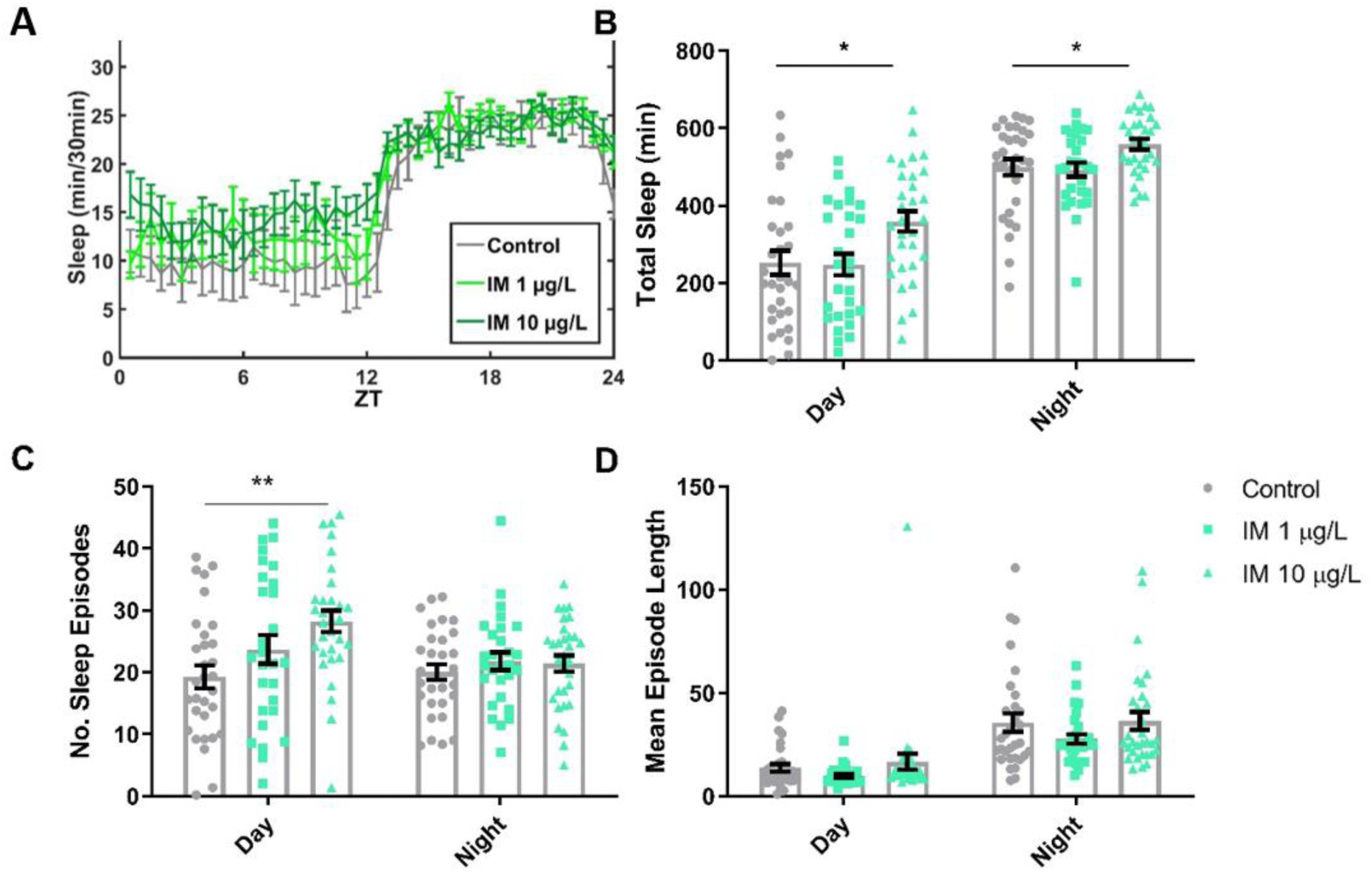
Field relevant concentrations of imidacloprid increases sleep in isolated bumblebee foragers. (A) Mean total sleep achieved for control *Bombus terrestris* foragers and those fed 1 µg/L (pale green line) or 10 µg/L (dark green line) imidacloprid (IM), per 30 min bin over the 24 h period. (B) Mean total sleep (min) for each treatment group in the day (*F*_2,87_=4.9, p=0.010) and the night (*F*_2,87_=4.1, p=0.019). (C) Mean number (No.) of sleep episodes initiated for each treatment group during the day (*F*_2,87_=5.4, p=0.006) and the night (*F*_2,87_=0.490, p=0.614). (D) Mean sleep episode length for each treatment group during the day (*F*_2,87_=1.7, p=0.182) and the night (*F*_2,87_=1.5, p=0.238). Each data point in the histograms represents a single bee, n=28-31 bees for each treatment.

*Bombus terrestris* foragers in a full colony setting showed diurnal rhythms in foraging activity (Figure 4-5). This rhythmicity was decreased in foragers exposed to 10 µg/L imidacloprid in both LD (Figure 4) and constant darkness (Figure 5). In LD conditions, imidacloprid decreased mean rhythmicity of foragers (Figure 4C) and increased the proportion of foragers who were arrhythmic from 48% to 65% (Figure 4B). Imidacloprid decreased foraging activity for both daytime and night-time (Figure 4D).

**Figure 4:**
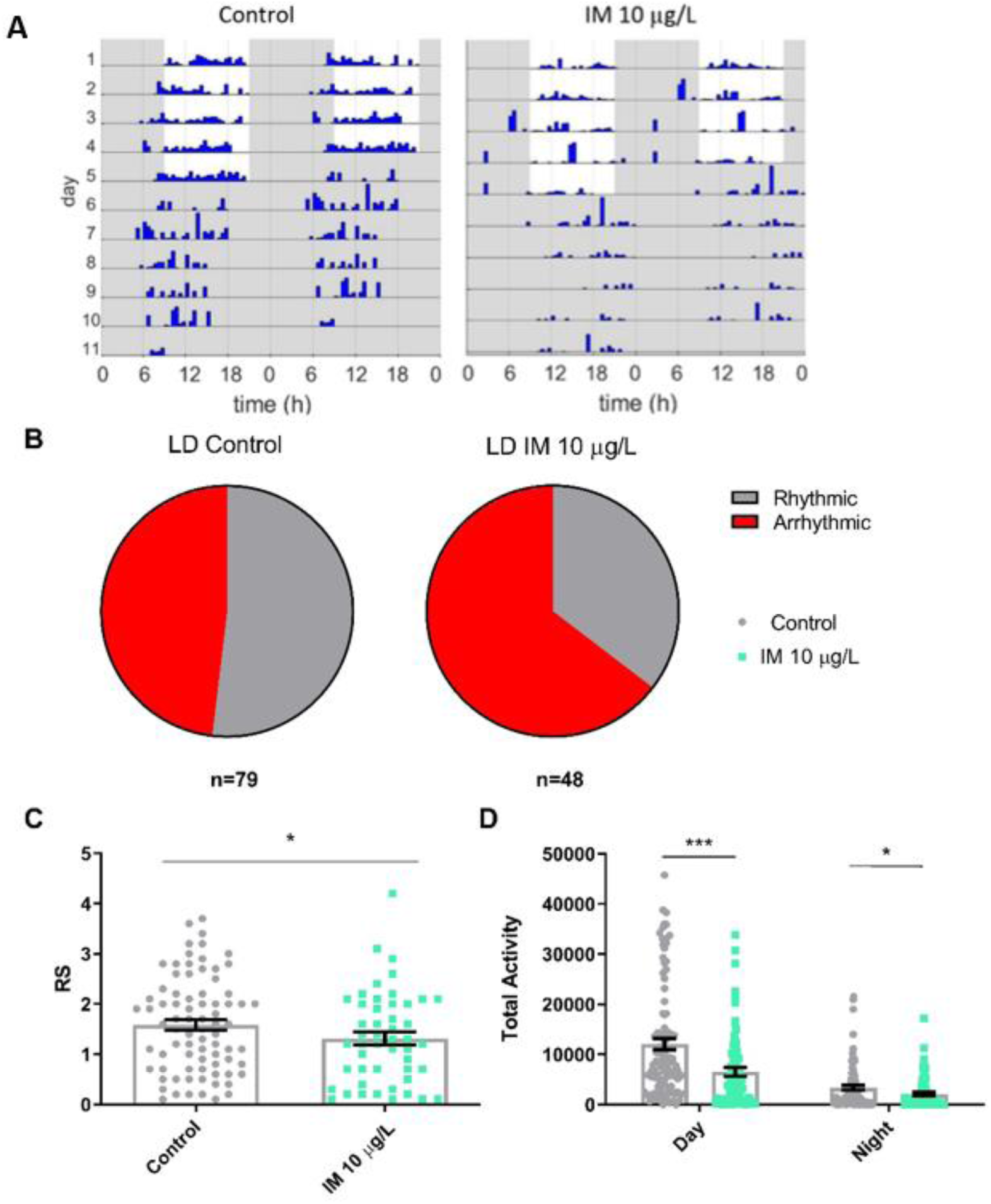
Field relevant concentrations of imidacloprid reduces foraging rhythmicity and activity in bumblebee foragers within the colony in 12 h:12 h light:dark conditions. (A) Representative actograms for a *Bombus terrestris* forager on control food and food concentration 10 µg/L imidacloprid (IM). (B) Proportion of foragers that were arrhythmic in LD for each treatment group. (C) Mean rhythmicity for either control foragers or foragers fed 10 µg/L IM in LD (*t*_125_=2.0, p=0.048). (D) Mean foraging activity for foragers in each treatment group in LD, during the day (*t*_170_=3.8, p<0.001) and the night (*t*_170_=2.0, p=0.042). Each data point in the histograms represents a single bee, n=48-79 bees for each treatment for rhythmicity, n=74-100 bees for each treatment for activity.

**Figure 5:**
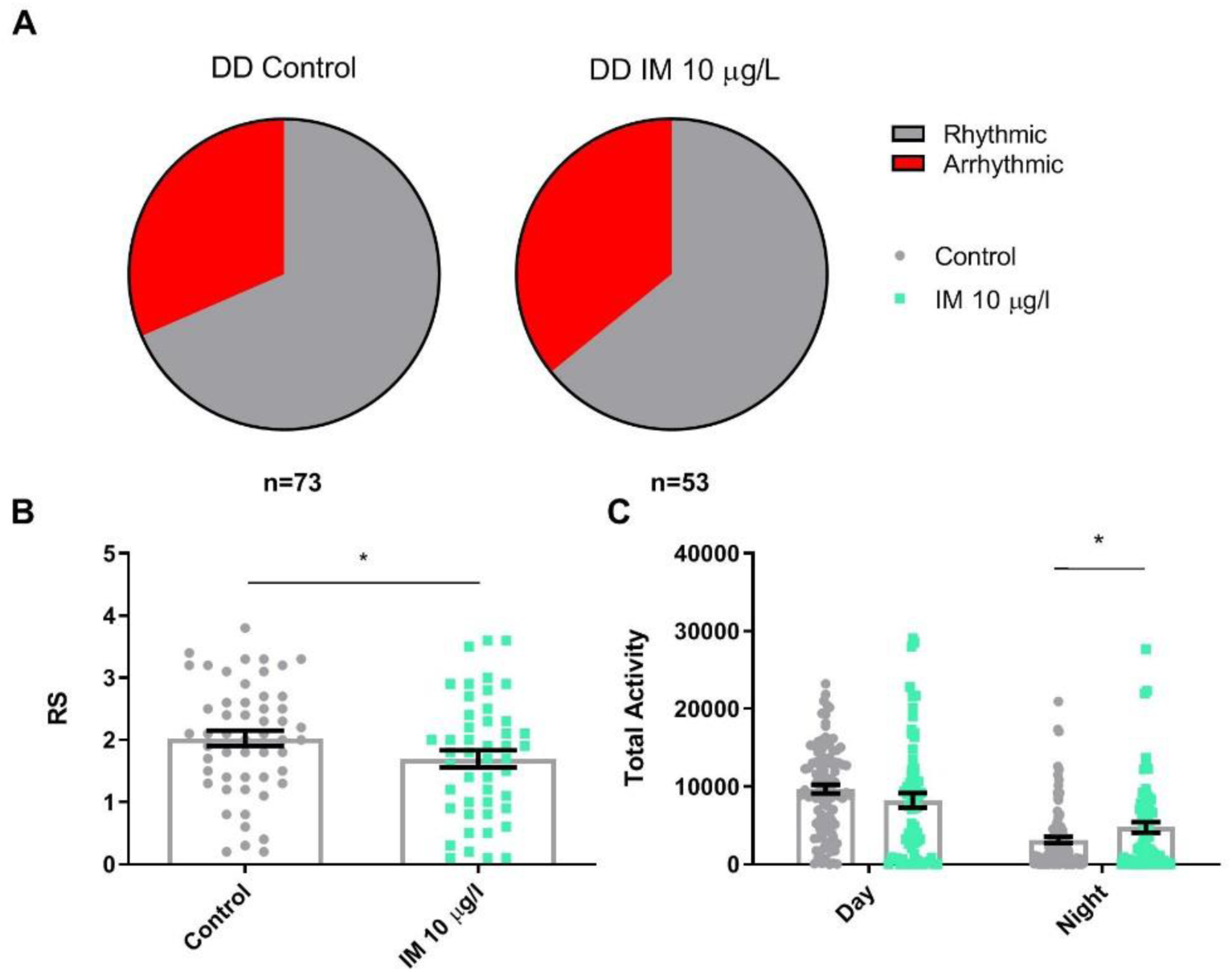
Imidacloprid reduces foraging rhythmicity and activity in bumblebee foragers within the colony in constant darkness. (A) Proportion of the *B. terrestris* foragers that were arrhythmic in DD on control food and food containing a field relevant concentration of 10 µg/L imidacloprid (IM). (B) Mean rhythmicity for foragers in each treatment group in DD conditions (*t*_125_=2.2, p=0.029). (C) Mean activity for foragers in each treatment group in DD conditions, during the subjective day (*t*_169_=1.6, p=0.105) and night (*t*_114_=-2.0, p=0.043). Each data point in the histograms represents a single bee, n=53-73 bees for each treatment for rhythmicity, n=74-100 bees for each treatment for activity.

In DD conditions, imidacloprid reduced mean rhythmicity for foragers (Figure 5B) and increased foraging activity during the subjective night (the 12h which were dark during the entrainment period) (Figure 5C). The proportion of foragers that were arrhythmic for control and imidacloprid exposed bees were similar in DD, 31% and 36% respectively (Figure 5A). Therefore, field relevant imidacloprid concentration disrupt circadian rhythmicity and increase mistimed daytime sleep in isolated *Bombus terrestris* foragers, and reduced foraging rhythmicity for foragers in the colony. Foragers both in isolation and the colony showed that neonicotinoids cause a profound disruption of timing of activity with reduced daytime activity.

## DISCUSSION

We show that imidacloprid causes a reduction in and mistiming of locomotion and foraging activity of bumblebees, increasing the proportion of foraging that occurs at night and increasing daytime sleep. Imidacloprid has previously been shown to reduce locomotion in isolated *Bombus terrestris*[39], sweat bees like *Melipona quadrifasciata*[40] and solitary bees like *Osmia bicornis*[41], and to reduce in-nest activity in another bumblebee *Bombus impatiens*[15]. We found that imidacloprid exposure also reduced the foraging activity of foragers within the colony, with individuals making fewer foraging trips in LD conditions. This is consistent with previous, field-based studies showing that exposure to imidacloprid under field conditions decreased the number of foraging trips carried out by foragers[42, 43], and neonicotinoids reduced the quantity of sucrose and pollen collected by *Bombus terrestris* colonies[44]. This could in part be driven by reduced foraging motivation caused by the apparent appetite suppression that imidacloprid can cause, with 10 µg/L shown to decrease feeding by 30% in *Bombus terrestris*[13].

We also show that field relevant doses of imidacloprid impact locomotor and foraging rhythmicity. Imidacloprid reduced the rhythmicity of daily activity in foragers in LD conditions, suggesting that the neonicotinoid may be interfering with light input into the clock. Light signalling from the visual circuit and the Hofbauer-Buchner (H-B) eyelet to the clock neurons is dependent on nAChRs so this is a potential route for disruption [4, 21]. Foraging rhythmicity in DD was also reduced, suggesting that imidacloprid may also disrupt the circadian clock, potentially through acting as an agonist at these nAChR synapses which are between the light sensing organs and the clock neurons. Previous work showed nAChR agonists can directly stimulate clock neurons in a cockroach *Rhyparobia maderae* causing increased calcium influx through voltage-gated calcium channels[3]. Likewise, nAChR agonists increase calcium influx of *Drosophila* PDF releasing lateral ventral neurons (LNvs)[45, 46]. Conversely nAChR antagonists block spike-dependent nAChR synaptic signalling required for rhythmic LNv activity[23]. Work in another cockroach, *Periplaneta americana*, showed neonicotinoids can also act on the thoracic ganglia, which control motor function in insects[47]. Thus, neonicotinoids may be acting directly through multiple neurons regulating both locomotion and circadian regulation of locomotion, causing activation and/or depolarisation block and hence compromising rhythmic foraging activity. In addition to direct classical pharmacological drug-receptor interactions, neonicotinoid exposure changes expression of hundreds of genes in worker bumblebees (*Bombus impatiens*), including genes which are involved in locomotion[47].

The reduction in rhythmicity observed for foragers in LD conditions and within the colony suggest that the effect of imidacloprid on rhythmicity cannot be mitigated by strong zeitgebers (*i*.*e*. time-givers or entrainment signals) such as light or social cues[48, 49]. This may imply that the reduction in foraging rhythmicity observed for bumblebee colonies exposed to imidacloprid in the lab are likely to reflect deleterious consequences in the field, as is the case for reductions in foraging activity[42, 43]. A disruption to the clock in foragers may further reduce their foraging efficiency as they will not be able to form the time-memories required to accurately visit different flowers[5, 50]. The clock also feeds into the sun-compass navigation pathway that foragers use to navigate[29]. Neonicotinoids have previously been shown to reduce homing ability in honeybees[51] and disruption to the clock could be a contributing factor to this. Reduced foraging efficiency is likely to reduce the capacity of the colony to grow and reproduce. Reduced feeding and foraging is associated with less brood production[52] and smaller colonies which are less resilient and less likely to produce queens[53].

Field relevant concentration of imidacloprid also disrupted sleep, with 10 µg/L increasing daytime sleep in foragers, hence the reduction in daytime locomotor and foraging activity observed. The mushroom body, which is known to regulate sleep/wake cycle in insects, contains groups of both wake and sleep promoting Kenyon cells and signal *via* nAChRs to the mushroom body output neurons which are also important for sleep regulation[25, 54, 55]. Sleep promoting neurons have been shown to be specifically activated by nAChRs[56], providing a possible route for imidacloprid to induce sleep. Furthermore, whole-cell patch clamp recordings from bumblebee Kenyon neurons of the mushroom body showed that they were directly stimulated by imidacloprid[57]. The increase in inappropriate daytime sleep observed may also further indicate a disruption to the circadian clock, as the timing of sleep is dictated by the clock[31].

Neonicotinoids reduce activity and foraging motivation in pollinators[43], in addition continued exposure causes bees to prefer neonicotinoid-laced food sources[58, 59], further exacerbating their negative impact. Here we demonstrate that very low field relevant concentrations of neonicotinoids disrupt rhythmicity of foraging activity as well as increasing daytime sleep, further reducing the opportunities for bees to forage and pollinate and having knock-on effects on circadian and sleep regulated physiological and behavioural processes in the bee. This is likely to have a detrimental effect on colony fitness in the field as well as reducing the yield of crops and wild plants reliant on bee pollination. Furthermore, we establish a number of highly sensitive (down to 1 ppb neonicotinoids) high throughput behavioural assays for measuring the detrimental sublethal effects of insecticides on pollinators.

## MATERIALS AND METHODS

Further information and requests for resources should be directed to Dr James JL Hodge.

### *Bombus terrestris* colonies

*Bombus terrestris audax* colonies (Biobest), containing cotton wool and approximately 80-100 workers, were ordered through Agralan (#BB121040-CF1). They were maintained at 21°C, 12h:12h LD, with a 30 mins dawn/dusk period where lights were at 50%. Colonies were provided with Biogluc® (Biobest) *ad lib* in the foraging arena and 1 teaspoon pollen (Agralan #BB008513) every 5 days which was introduced into the nest box. Imidacloprid (PESTANAL Sigma-Aldrich #37894), was administered via the Biogluc®, at field relevant concentrations of either 1 or 10 µg/L[23]. Colonies were attached via a clear plastic tube (15 mm diameter and 200 mm length), to a foraging arena (1000 × 50 × 50 mm) purpose built by the University of Bristol mechanical workshop out of UV-transmitting acrylic. A wide ramp from the entrance to the foraging arena to the floor of the arena was built from card and duct tape to ensure that foragers could return to the nest box in darkness, when they cannot fly[60].

### Circadian rhythm analysis for isolated foragers

For the collection of circadian and sleep data for bumblebees, a system was set up using the Locomotor Activity Monitor (LAM16, TriKinetics Inc, USA) [61]. Foragers were collected from the foraging arena and loaded into tubes (PGT16×100). At one end of the tube was a rubber cap (CAP16-Black) with a 1 g silica packet glued inside to control the humidity. At the other end, a 15 mL Falcon tube with a small whole drilled near the base was attached. This contained Biogluc® with or without neonicotinoids, allowing bees to feed *ad libitum* throughout the experiment. The tubes were loaded into the LAM and monitored for five days in LD and five days in DD conditions. In the monitor an infrared beam crossed the tube and every beam break was counted in real time by a host computer. Each beam break was counted as a single activity bout, allowing the total activity for each bee to be summed per day and per 30 mins and 1 min bins for circadian analysis. For each bee, the rhythmicity statistic (RS), period length and total daily activity levels were then calculated. Circadian analysis was carried out using *Flytoolbox*[83] in MATLAB (MATLAB and Statistics Toolbox Release 2015b). Conventionally, an RS >2 is rhythmic, an RS of 1.5-2 is weakly rhythmic and an RS ≤1.5 is arrhythmic[38]. The day and night activity levels for each bee were calculated using the *daynight* program[62] in MATLAB. The individual bee’s period length was used to split the activity data into subjective days and nights and then the activity counts for each were summed. The mean daily activity in daytime and night-time for the five days was then calculated.

### Sleep analysis for isolated foragers

Sleep is defined as any inactivity lasting more than five minutes, as has been done in previous studies[33]. Sleep analysis was carried out on the five days of LD[63], using activity data that had been summed into both 1 min and 30 mins bins. Analysis was performed using the Sleep and Circadian Analysis MATLAB Program (SCAMP[64]). The mean total quantity of sleep per 30 mins bin was calculated and displayed. Total quantity of sleep for the day and night for each bee was quantified and the mean taken for the five days. Other sleep measures reported were the number of sleep episodes initiated during the day and night and the mean length of these episodes.

### Foraging rhythmicity assay for bumblebees in the colony

Circadian rhythmicity within the colony was assayed using a micro radio frequency identification (RFID) setup[65]. Approximately 40 foragers were collected from the foraging arena. These were anaesthetised using CO_2_ and an RFID tag (Microsensys GmbH mic3-TAG) was stuck to the centre of their thorax with superglue (Loctite). Foragers were then returned to the colony nest box. After a day for acclimatisation and recovery from the CO_2_ exposure, recording began. An RFID reader (Microsensys GmbH iID®MAJAreadermodule4.1) was placed at the entrance to the foraging arena so that foragers had to pass through the reader to enter the arena. The data from the readers were collected on a host (microsensys GmbH iID®HOSTtypeMAJA4.1), for a 10 days period; 5 days LD and 5 days DD, as in the LAM experiments above. The data were then summed into 30 mins bins, with each pass through the reader being counted as a single activity bout. These data were then analysed as above using the *Flytoolbox* in Matlab. This was repeated with three colonies for each treatment group. Pollen was provided the day before recording began and on day 5. Biogluc was available *ad libitum* in the foraging arena and was either untreated for control groups or contained 10 µg/L imidacloprid for treated groups. Three colonies were tested for each treatment group.

### Statistical Analysis

Analysis was carried out using a one-way ANOVA and the results displayed in figure legends. First the data was checked for normality using a Shapiro-Wilk test. The homogeneity of variance was also tested using Levene’s test for equality of variances. Means were then compared using a one-way ANOVA with *post hoc* pairwise comparisons using Tukey’s multiple comparisons test. Statistical analysis was done in IBM SPSS Statistics 24. Graphs were created in GraphPad (Prism version 8.0.0). For all histograms, every data point was plotted with lines showing the mean ± standard error of the mean (SEM). Where *post hoc* tests were done, these were displayed as *p* ≤ 0.05*, *p* ≤ 0.01**, *p* ≤ 0.001***, *p* ≤ 0.0001****).

### Data and Software availability

The datasets generated during this study are available at [name of repository] [accession code/web link].

## Acknowledgements

We thank Drs Stephen Montgomery, Edgar Buhl and Herman Wijnen for providing useful comments on the manuscript. This work was supported by a BBSRC studentship BB/J014400/1 and Leverhulme project grant RPG-2016-318 awarded to JJLH.

## Author Contributions

Circadian and sleep assays were carried out by KT. KT wrote the first draft of the paper. The project was supervised by JJLH and SAR, who secured funding and edited the manuscript.

